# Oxygen is Toxic in the Cold in *C. elegans*

**DOI:** 10.1101/2024.07.17.603990

**Authors:** Cameron M. Suraci, Michael L. Morrison, Mark B. Roth

## Abstract

Temperature and oxygen are two factors that profoundly affect survival limits of animals; too much or too little of either is lethal. However, both humans and animals can exhibit exceptional survival when oxygen and temperature are simultaneously low. To better understand this apparent synergistic interaction between oxygen and temperature, we assayed the survival of *Caenorhabditis elegans* in experimental environments at different temperatures and oxygen concentrations. While nematodes cannot survive a day in room air at 2 °C (cold shock), we found that when oxygen is low, 200-fold less than room air, they can survive this temperature for 48 hours. Consistent with this, we found that worms exposed to high oxygen concentrations, 35 times greater than room air, are more sensitive to low temperature than worms in room air. These results show that normal atmospheric levels of oxygen are toxic in the cold. Using these survival assays, we found that cold acclimatization protects worms from lethal effects of high oxygen, mutations in the cold acclimatization pathway affect oxygen tolerance, and that the naturally occurring and physiologically relevant compounds glucose, manganese (II), and ascorbate improve survival limits in both low temperature and high oxygen when supplied early in life. These results show that the interdependence of temperature and oxygen on survival in *C. elegans* is based in part on shared physiological mechanisms involved in response to these two environmental stressors. The evolution and natural history of stress responses in animals suggest similar phenomena may function in humans.

## Introduction

Prolonged exposure to low oxygen or low temperature is lethal in most animals. Despite this, some humans, such as airplane wheel-well stowaways, have anomalously survived conditions where either the low oxygen alone or the cold alone would have been enough to ensure death^1^. One possible explanation of this anomaly is that low oxygen and low temperature are synergistically protective. The protection the cold provides against the effects of low oxygen has been applied in humans: cooling is used to prevent brain damage in hypoxic infants^2^. However, the protection that low oxygen provides against the effects of the cold is less understood and has not been applied in a therapeutic setting to our knowledge.

To understand more about the relationship between oxygen and temperature we have been studying the nematode *C. elegans*. This animal has been shown to be sensitive to both low oxygen^3^ and cold shock (2 °C)^4^; additionally, worms raised at 12 °C in room air and then challenged with cold shock in room air survive^5^. Toxicity of excess oxygen, however, has not been previously demonstrated in *C. elegans*, as they are capable of surviving and reproducing for up to 50 generations in 100% oxygen at atmospheric pressure^6^. Here, we use changes in temperature and oxygen to show that nematode death in the cold is due to oxygen toxicity. We go on to show that the responses to high oxygen and low temperature are highly overlapping, as the same mutants and naturally occurring compounds affect tolerance to each stress.

## Results

### Normal Oxygen Levels are Toxic in the Cold

Previously, it has been shown that cold (2 °C cold shock)^4^ and anoxic (100% nitrogen) conditions^3^ can be lethal to adult *C. elegans*; our data shows that these conditions kill nearly 100% of animals in 24 and 48 hours, respectively. Here, we show that combining these two conditions results in a higher survival rate than either one alone. Fig. 1a shows that most first-day adults in 23 °C nitrogen or 2 °C room air died after 48 hours, while adults in 2 °C nitrogen had an average survival rate of 93%. To further explore this phenomenon, we titrated the oxygen concentration in a 24-hour, 2 °C cold shock (Fig. 1b). On average, 98% of the nematodes exposed to 0.10 kPa of oxygen survived the cold shock. An increase to just 0.50 kPa oxygen (40 times less than room air), however, was nearly 100% lethal.

**Fig. 1:**
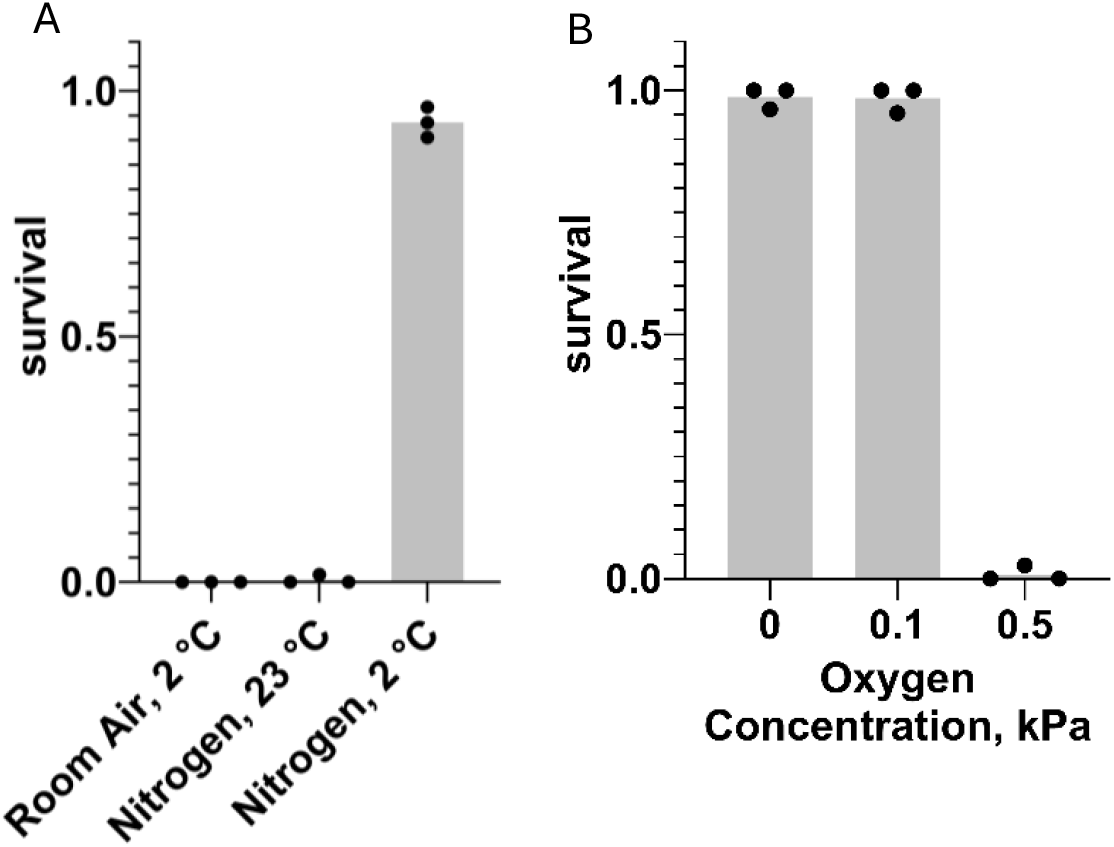
Effect of Oxygen Concentration on *C. elegans* Cold Shock Survival. Each point represents the survival of a plate of 20 to 100 nematodes. **A**. Survival after 48 hours in the conditions described on the X axis. **B**. Survival after a 24-hour 2 °C cold shock in nitrogen containing low oxygen concentrations.

### High Oxygen is Toxic at Room Temperature

*C. elegans* can complete their life cycle in 100% oxygen at room pressure^6^. We found that slightly increasing the pressure of oxygen to 136 kPa does not kill egg-laying adults, but it does prevent their progeny from developing to adults. Further increasing the pressure of oxygen to 377 kPa or 687 kPa (687 kPa of oxygen is hereafter referred to as hyperbaric oxygen/HBO) causes mortality in the first 8 hours of exposure (Fig. 2). Increasing oxygen pressure was associated with a decrease in survival at all time-points after the first two hours. Additionally, the survival rate of nematodes in 687 kPa of oxygen negatively correlated with length of exposure at all time-points. Importantly, the mortality observed in these assays is dependent on the partial pressure of oxygen, not total pressure: when egg-laying adults were exposed to a mixture of 99.5% nitrogen and 0.5% oxygen gas raised to 1103 kPa (with an oxygen partial pressure of 5.5 kPa), no mortality was observed during their egg laying period and their progeny were able to grow to adulthood. These results show that oxygen at high concentration is a dose and time-dependent toxin in *C. elegans*.

**Fig. 2:**
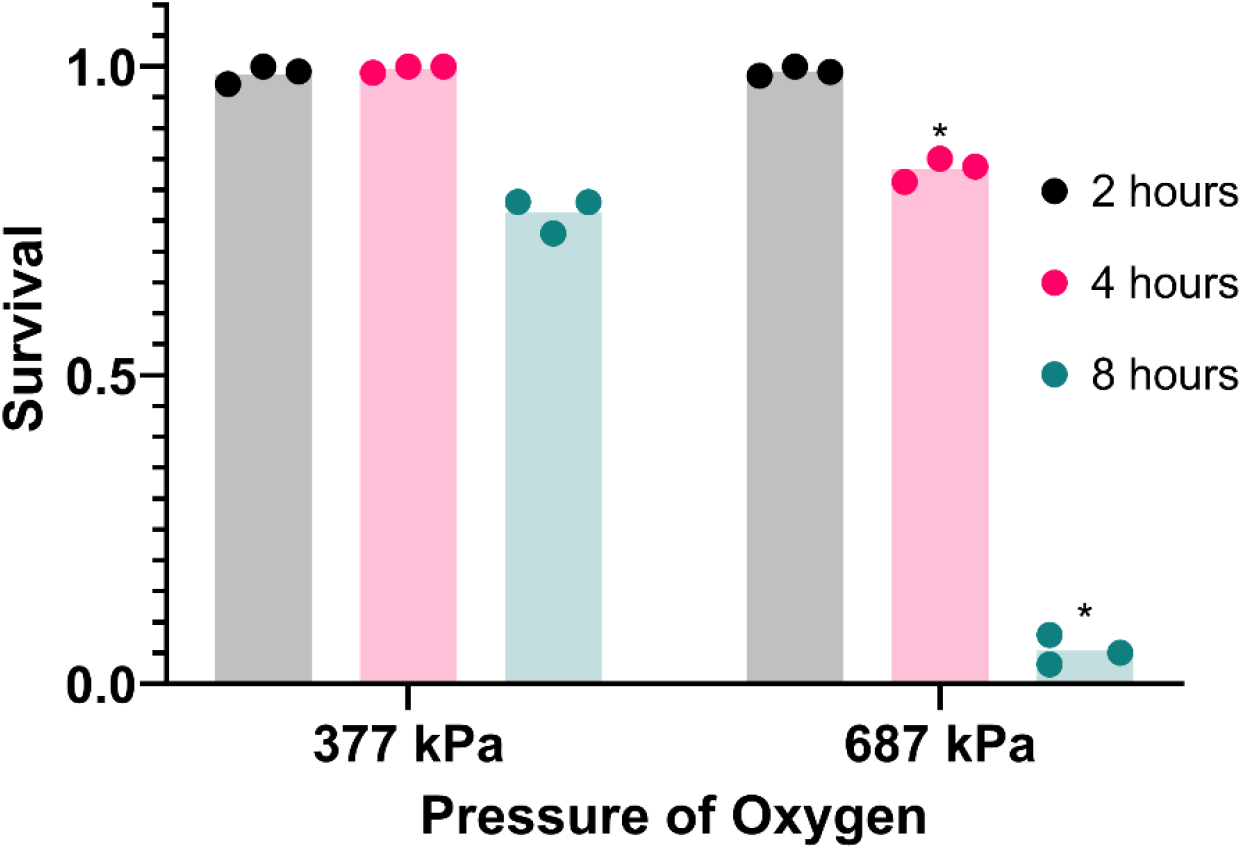
Hyperbaric oxygen (HBO) is toxic. Survival of *C. elegans* in high oxygen. Each point represents the survival of a plate of 20 to 100 nematodes. *P < 0.05 when comparing to 377 kPa.

### Low Temperature Increases the Toxicity of Hyperbaric Oxygen

Here, we show that HBO and cold shock synergize to cause greater lethality than either condition alone (Fig. 3). Nematodes exposed to room temperature HBO or cold room air for two hours had average survival rates of 95% and 93%, respectively, whereas nematodes exposed to cold HBO for the same amount of time had an average survival rate of just 31%, an almost three-fold decrease from the expected survival rate of 88% if these conditions acted independently of one another.

**Fig. 3:**
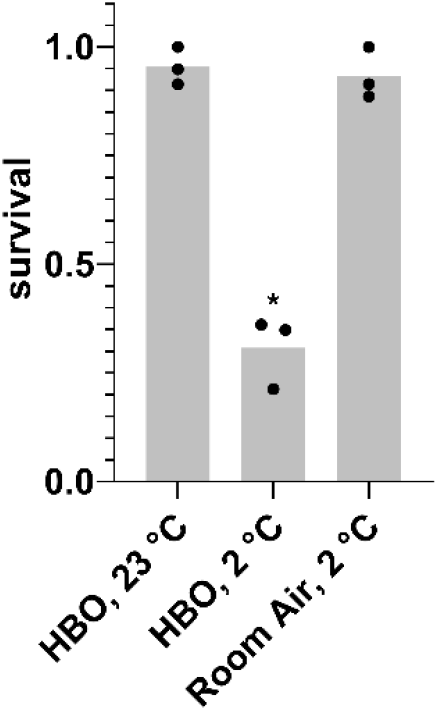
Effect of 2-hour exposure to combined cold shock and HBO. Survival of *C. elegans* after a 2-hour exposure to room temperature HBO, 2 °C HBO, or 2 °C room air. Each point represents the survival of a plate of 20 to 100 nematodes. *P < 0.05 when 2 °C HBO is compared to either 23 °C HBO or 2 °C room air.

### Cold Acclimatization Confers Resistance to Hyperbaric Oxygen

If oxygen is toxic in the cold, then prior acclimatization to the cold, which increases survival during cold shock^5^, may also confer resistance to HBO. Consistent with this prediction, we show that prior acclimatization of nematodes by growing them at 12 °C dramatically increases their capacity for survival in room temperature HBO compared to those grown at room temperature, effectively tripling the LD^50^ from 4 hours to 12 hours (Fig. 4).

**Fig. 4:**
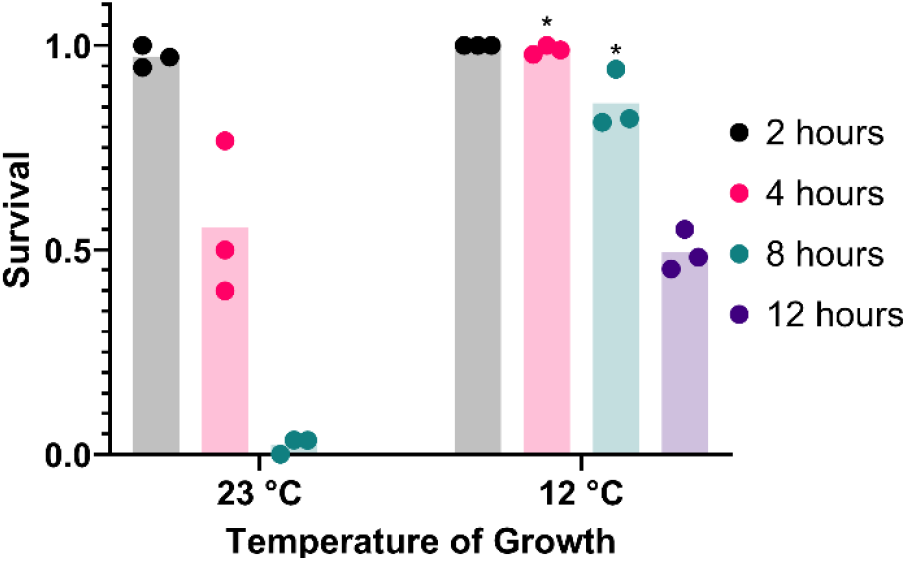
Cold acclimatization provides resistance to hyperbaric oxygen. Survival of animals grown from L1 stage at 23 °C or 12 °C and subsequently exposed to HBO as adults. Each point represents the survival of a plate of 20 to 100 nematodes. *P < 0.05 when comparing to animals grown at 23 °C.

### Strains Exhibiting Sensitivity and Resistance to Both Cold Shock and Hyperbaric Oxygen

If oxygen is toxic in the cold, then mutations that affect cold tolerance may also affect HBO tolerance. It has been previously shown that age-1(-), unc-104(-), and akt-1(-) strains of *C. elegans* have enhanced resistance to cold shock compared to wild type^7^, while deg-1(-) strains are cold acclimatization defective^8^. Here, we tested if these mutants also have atypical survival in HBO at room temperature (Fig. 5). We found that TJ1052 (age-1), CB1265 (unc-104), and BQ1 (akt-1) strains exhibited enhanced survival rates in HBO relative to wild type, while the strain TU38 (deg-1) had a decreased survival rate, demonstrating an overlap between cold tolerance and oxygen tolerance pathways.

**Fig. 5:**
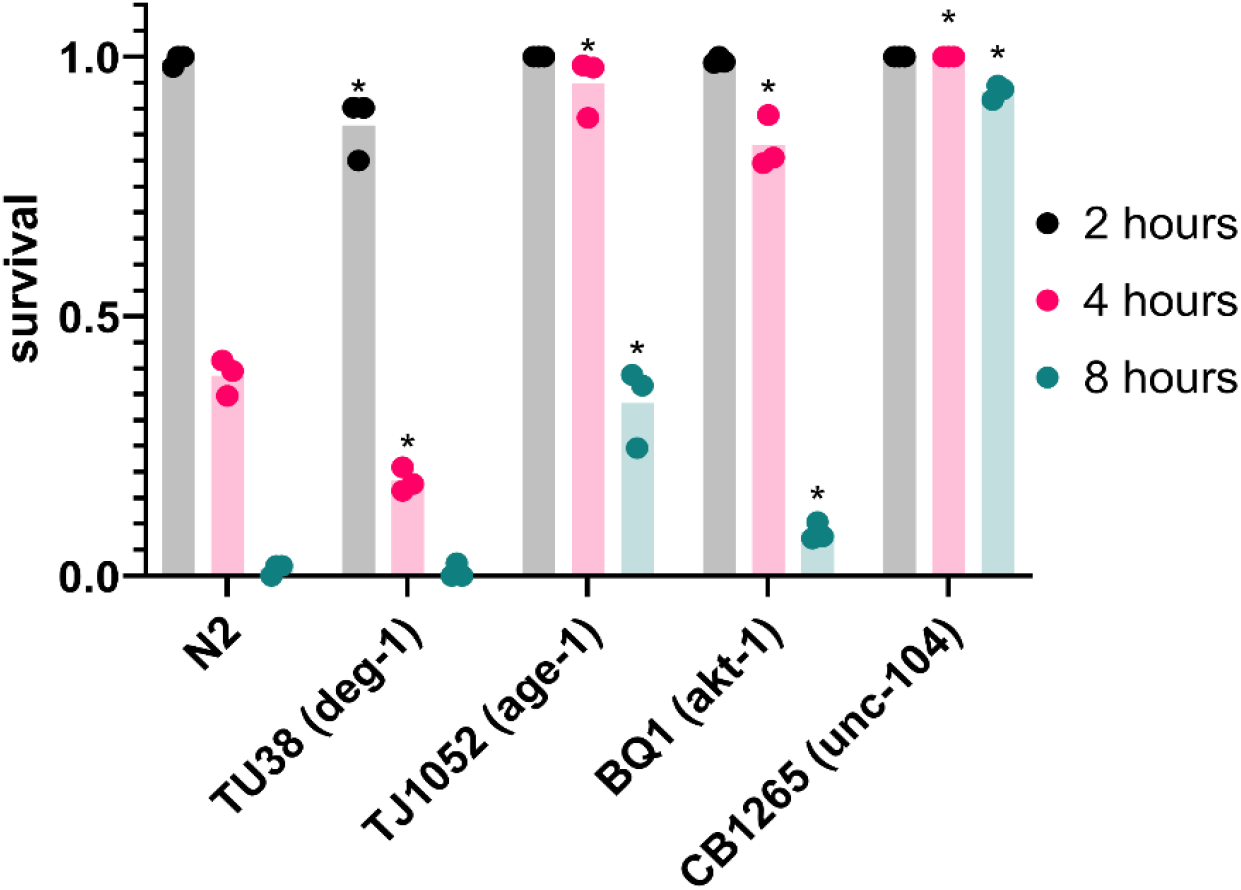
Genetic background effects survival in hyperbaric oxygen. Survival of mutant strains in HBO. Each point represents the survival of a plate of 20 to 100 nematodes. *P < 0.05 when comparing to wild type (N2).

### Diverse Compounds Confer Resistance to Both Cold Shock and Hyperbaric Oxygen

If oxygen is toxic in the cold, then compounds that confer resistance to cold shock may also confer resistance to HBO. Here, we demonstrate that glucose, manganese (II) chloride, and ascorbate each enhance the survival of *C. elegans* in both cold shock and HBO (Fig. 6). Nematodes were grown from the L1 stage to adulthood on plates containing either 300 mM glucose, 3 mM MnCl^2^, 100 mM ascorbate, or no additive, before being exposed to either HBO or cold shock. All three additives induced increased survival at most timepoints in both conditions. These results provide further evidence that cold tolerance and oxygen tolerance are overlapping pathways.

**Fig. 6:**
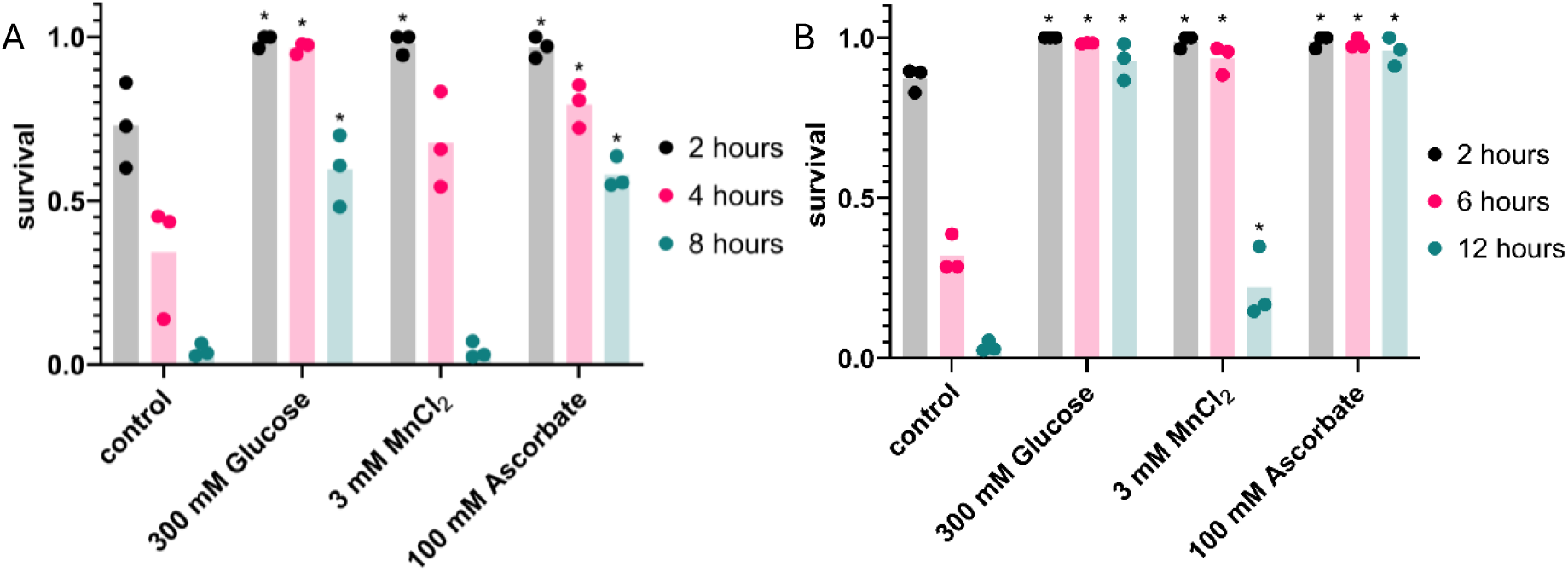
Glucose, MnCl_2_, and ascorbate increase both cold shock and hyperbaric oxygen resistance. Survival in 2 °C cold shock (**A**), or HBO (**B**), after animals were grown with compounds listed on the X axis. Each point represents the survival of a plate of 20 to 100 nematodes. *P < 0.05 when comparing to control untreated animals.

## Discussion

In this research, we showed significant overlap between the pathologies of cold shock and oxygen toxicity in *C. elegans*. We showed that nematodes survived cold anoxia better than room temperature anoxia or cold room air. We demonstrated that HBO could kill nematodes, and that low temperature increased the toxicity of HBO. We also showed that we could improve HBO survival by cold-acclimatizing nematodes, and that mutations to genes in the cold acclimatization pathway affected HBO tolerance. Finally, we identified naturally occurring and physiologically relevant compounds that improved survival limits in both cold shock and HBO. These results suggest that oxygen toxicity is increased in the cold and contributes to cold-related death in *C. elegans*.

The increased toxicity of oxygen in the cold is likely the result of a variety of factors. Decreasing the temperature from 23 °C to 2 °C increases the solubility of oxygen in water by approximately 57%, which is not enough to induce oxygen toxicity on its own; a 5-fold increase above atmospheric oxygen concentrations was necessary to induce any negative effects in nematodes at room temperature. However, other processes may account for the toxicity we observed. The rate- reduction of temperature-sensitive enzymes responsible for the neutralization of reactive oxygen species (ROS) and control of redox potential may limit the ability to prevent oxidative damage.

Notably, *C. elegans* decrease oxygen consumption as temperature decreases^9^, which may be an adaptation to reduce the potential for oxidative damage; oxidative metabolism has been implicated as a way that organisms produce ROS from oxygen^10^. Additionally, if oxygen-dependent processes proceed while temperature-sensitive ones are slowed or stopped, the organism may proceed into non-functional states. Our lab has previously shown that this occurs in cold-shocked 2-cell *C. elegans* embryos; these embryos were incapable of cytokinesis in the cold but would continue to produce up to 8 centrioles in each blastomere, resulting in failure of chromosome segregation, aneuploidy, and death upon restoration of the cell cycle when returned to room temperature^11^. Embryos that were anoxic during the cold exposure did not produce excess centrioles and, as a result, were viable upon restoration of room air and temperature. The difference in cold shock survival rates between nematodes exposed to 0.1 and 0.5 kPa oxygen is further indication that limiting oxidative activity is beneficial in the cold; mammalian mitochondria are only capable of functioning at oxygen concentrations above 0.5 kPa, below which they will initiate cellular apoptosis in normothermia^12^. A model to explain the synergy between anoxia and cold is that lowering the oxygen concentration to the point that mitochondria do not function may prevent oxidative activities that are toxic in the cold; in turn, the low temperature may slow the use of stored sugars, extending anerobic survival.

The resistance to HBO shown by *C. elegans* acclimatized to cold suggests that animals in the wild have sufficient time during temperature change to enact changes that protect them from oxidative stress. In our research, we focused on genes related to cold acclimatization regulation to emphasize the relationship between activation of this pathway and oxygen toxicity resistance; future research could focus on the specific genes affected by this pathway that confer this resistance. ASJ neurons use insulin to prevent intestinal neurons from initiating a cold-tolerance response; as the temperature drops, the ASJ neurons produce less insulin^7^. These neurons are incapable of transmitting this signal in worms without unc-104 (a kinesin), while age-1(-) and akt-1(-) worms have intestinal neurons that are insensitive to the insulin signal. Notably, age-1(-) nematodes have a lower metabolic rate than wild type^9^, a phenotype which may confer resistance to both cold shock and HBO. The deg-1 gene product is a sodium channel that acts as a thermoreceptor in ASG neurons, which are responsible for positive regulation of cold tolerance^8^. Our results show that this gene is also necessary for stress response to high oxygen. Scnn1a, the rat ortholog of deg-1^13^, shows increased gene expression and protein activity in the alveolar cells of hyperoxic rats^14^. If deg-1 activity is similarly affected by high oxygen, it could explain this gene’s importance to HBO tolerance. Further research is necessary to determine if oxygen concentration affects deg-1.

The finding that glucose, manganese (II), and ascorbate can protect nematodes from HBO and cold shock provides further evidence that the pathologies of oxygen and cold toxicity are highly overlapping. Ascorbate and manganese (II) have both been implicated in the neutralization of ROS, a potential source of HBO and cold shock toxicity. Ascorbate acts as a direct antioxidant against ROS^15^, as well as acting as a cofactor in a variety of enzymes^16^ necessary for ROS-reducing activities, such as carnitine production^17^. Manganese (II) is used by many *Lactobacillus* strains in place of superoxide dismutase^18^ and acts as a catalyst for hydrogen peroxide dismutation in the presence of biological levels of bicarbonate^19^. The mechanism by which glucose prevents HBO and cold shock toxicity may be related to the intracellular oxidation state; in human cells, it is known that hyperglycemia^20^ and anoxia^21^ increase the NADH/NAD+ ratio, while hypothermia decreases it^22^. The highly oxidizing environment of HBO likely reduces this ratio as well. The reductive pressure produced by glucose supplementation may counteract the oxidative environments produced by HBO and cold shock, resulting in the enhanced tolerance to these environments that we observed. Likewise, research has shown that combining two oxidative conditions (HBO and cold shock, fig.3) or two reductive conditions (hyperglycemia and anoxia)^23^ results in decreased survival of nematodes. Future research is necessary to confirm the effects of these environments on the intracellular oxidation state.

Is oxygen toxicity in the cold a nematode-specific phenomenon, or is it found in other animals? Animals ranging from protozoans to mammals are known to decrease their body temperature in hypoxic conditions^24^. While it is not evidence of oxygen toxicity in the cold, this adaptation mirrors our result regarding survival of nematodes in cold nitrogen. Research has also shown that humans, similar to *C. elegans*, decrease their whole-body oxygen consumption by half for every 10 °C reduction in temperature, a phenomenon known as the Q10 effect^25^. While the use of anoxia to treat hypothermia has not been attempted to our knowledge, recent analysis has shown that hypothermia patients with hyperoxic arterial oxygen concentrations had a 28-day mortality rate twice that of normoxic patients^26^. Further research to understand if the observations made here extend to other animals and humans may improve our ability to use oxygen concentration and temperature to improve clinical outcomes.

## Methods

### Plates and Bacteria

Due to the uncontrollable variation in composition of peptone and agar, peptone-free agarose-based plates were used. For 1 liter of this solution, 3 g of NaCl and 15 g of agarose was added to 975 mL of agarose and autoclaved for 1 hour. After cooling, 1 mL of cholesterol (5 mg/mL in ethanol), 1 mL of 1 M CaCl_2_, 1 mL of 1 M MgSO_4_, 1 mL of uracil (2 mg/mL in water), and 25 mL of 1 M (pH 6.0) KPO_4_ buffer were added. 35 mm and 60 mm plates were filled with 5 mL and 15 mL of solution, respectively, and stored at 4°C. 35 mm plates with additional additives were created by adding concentrated solutions of the additive to the molten agarose. It was necessary to adjust the ascorbate solution to a neutral pH to stay within the KPO_4_ buffering capacity.

*E. coli* (OP50) was grown in 1 liter of standard LB medium overnight at 37 °C on a shaker running at 200 rpm. The bacteria were collected by centrifugation at 4000 rpm for 15 minutes, before resuspension in 40 mL of autoclaved water. This solution was added to 60 mm plates as necessary to maintain worm populations. For 35 mm plates, 100 µL of a 10x dilution of this solution was added to the center of each plate.

### Nematode Maintenance

Bristol N2, TJ1052 (age-1), CB1265 (unc-104), TU38 (deg-1), and BQ1 (akt-1) strains were sourced from the Caenorhabditis Genetics Center. Nematodes were grown on 60 mm plates from eggs to adulthood. To produce a new synchronized generation, adults were minced using a razor blade and the released eggs were distributed onto a new plate. Room temperature raised-worms were grown on the bench for 3 days to reach adulthood, while 12 °C-raised worms were grown in an incubator for 9 days to reach adulthood. For survival assays, these populations were then washed off the 60 mm plates using water. The adults were then settled and resuspended in water 3 times to remove residual bacteria and juvenile nematodes. From this solution, 20 to 100 adults were transferred to 35 mm plates, which had any dead or juvenile nematodes removed from them after drying. When working with additives, nematodes were instead transferred to the control and additive plates as L1s, with dead and juvenile nematodes being removed once the populations had reached adulthood. CB1265 (unc-104) adults were transferred to plates using a platinum wire, as their uncoordinated phenotype prevented them from swimming; this made separating adults from juveniles using the described settling method inefficient, as they settled at similar rates.

### Cold Shock Assays

35 mm nematode plates were placed in sealed containers (approximately 500 cc in size) with two openings, allowing gas (either room air, 100% nitrogen, or nitrogen with 1000 or 5000 ppm oxygen) to be run through each one at 200 cc/min using a mass flow controller (Sierra Inst. Monterey, CA). Gases were humidified with a bubble chamber prior to reaching the nematodes. These sealed containers were placed into a 2 °C incubator. Nematode plates were moved to the benchtop after a time course was completed. Survival was assessed the day after removal from the condition. Nematodes in the nitrogen-room temperature condition were not placed into an incubator; the remainder of the procedure was identical. Instead of using the continuous-flow containers, nematodes in cold HBO were pressurized in a pressure chamber (see “Hyperbaric Oxygen Assays” section of Methods), which was then placed in a 2 °C incubator.

### Hyperbaric Oxygen Assays

Nematode plates were placed into a pressure chamber (Alloy Products, Waukesha, WI). After sealing, the chamber was flushed with 100% oxygen or 99.5:0.5 N_2_:O_2_ for 90 seconds by holding the pressure release valve open. The pressure release valve was then closed, and the chambers were pressurized. Chambers were pressurized to 687 kPa in hyperbaric oxygen assays unless otherwise specified. Due to the slight expansion of the pressure chambers caused by the pressurized gas, it was necessary to add additional gas 1-3 minutes after initial pressurization to restore the desired pressure; pressure was consistent after this. When a time course was completed, the chamber was depressurized to return the animals to room pressure/room air.

